# Distinct structure of cortical population activity on fast and infraslow timescales

**DOI:** 10.1101/395251

**Authors:** Michael Okun, Nicholas A. Steinmetz, Armin Lak, Martynas Dervinis, Kenneth D. Harris

## Abstract

Cortical activity is organised across multiple spatial and temporal scales. Most research on the dynamics of neuronal spiking is concerned with timescales of 1 ms − 1 s, and little is known about spiking dynamics on timescales of tens of seconds and minutes. Here, we used frequency domain analyses to study the structure of individual neurons’ spiking activity and its coupling to local population rate and to arousal level across frequencies ranging from 0.01 to 100 Hz. In mouse medial prefrontal cortex (mPFC), the spiking dynamics of individual neurons could be quantitatively captured by a combination of interspike interval and firing rate power spectrum distributions. The relative strength of coherence with local population often differed across timescales: a neuron strongly coupled to population rate on fast timescales could be weakly coupled on slow timescales, and vice versa. On slow but not fast timescales, a substantial proportion of neurons showed firing anti-correlated with the population. Infraslow firing rate changes were largely determined by arousal rather than by local factors, which could explain the timescale dependence of population coupling strength of individual neurons. These observations demonstrate how individual neurons simultaneously partake in fast local dynamics, and slow brain-wide dynamics, extending our understanding of infraslow cortical activity beyond the mesoscale resolution of fMRI studies.

## Introduction

A single action potential lasts about a millisecond, and a second suffices for a vast range of sensory-motor and cognitive behaviours, such as recognising pictures and sounds, getting up or sitting down, or recalling a memory. Accordingly, most neurophysiological research has focused on sub-second timescales. However, several neural processes occur over much longer timescales (Huk et al., 2018). Transitions between sleep and wakefulness and between different stages of sleep occur on timescales of minutes and hours (Weber and Dan, 2016; Lecci et al., 2017; Meisel et al., 2017). During wakefulness, changes in arousal can span tens of seconds and minutes, yet they affect performance in sub-second behavioural tasks (Harris and Thiele, 2011; Palva and Palva, 2012; McGinley et al., 2015). Dynamics on slower timescales has been revealed by resting-state fMRI (Raichle, 2015), which infers neural activity from the (slow) changes in blood supply to different areas of the brain. However, fMRI monitoring of neural activity is limited to the so called infraslow range of 0.01-1 Hz. Furthermore, both fMRI and other approaches to study mesoscale infraslow cortical dynamics - such as electro-and magneto-encephalography (EEG, ECoG, LFP, MEG, e.g. see Popa et al., 2009; Palva and Palva, 2012; Mitra et al., 2018), and intrinsic and voltage-sensitive fluorescent-protein or dye imaging in experimental animals (White et al., 2011; Chan et al., 2015; Kraft et al., 2017) - cannot characterise individual neurons’ relationships to infraslow activity.

The relationship of individual neurons to infraslow brain dynamics, and the relationship between a neuron’s coupling to infraslow and fast dynamics, is thus poorly understood. For example, how much can the firing rate of an individual neuron change over tens of seconds and minutes, and how can these slow dynamics be summarised quantitatively? To what extent are slow changes in firing rate correlated among neurons, and what is the structure of these slow correlations? To what extent are a neuron’s relationships to slow and fast firing rate fluctuations similar, and might they be driven by the same underlying mechanisms?

Here we addressed these questions by analysing multi-hour recordings of neuronal populations in mouse medial prefrontal cortex (mPFC), performed using chronically implanted high-density silicon probes. We found that neuronal spike trains have ***1/f*** power spectral density (PSD), and that PSD in combination with interspike interval (ISI) distribution suffices for an accurate quantitative model of single neuron spiking dynamics on both fast and slow timescales. Coupling between individual neurons and the population was timescale-dependent, with many neurons strongly coupled to population rate on fast timescales but weakly coupled on slow timescales, or vice versa. Furthermore, on slow but not fast timescales, neurons’ phase preference with respect to the population rate was bimodal. Finally, in frequencies ≤ 0.1 Hz population rate was highly correlated with arousal as reflected by the pupil area. These results suggest that dynamics on fast and infraslow timescales are distinct processes, and likely regulated by distinct mechanisms at the single neuron level.

## Results

To examine the intrinsic spiking dynamics of single cortical neurons on timescales extending to tens of seconds and minutes we used chronically implanted multisite silicon probes (Okun et al., 2016; Jun et al., 2017) to record the activity of neuronal populations in the frontal cortex. The recordings lasted 1.5-3 h in head-fixed mice, standing or sitting in a plastic tube, and 4-8 h in freely behaving mice residing in their home cage. The recordings were performed with 16-and 32-channel Neuronexus probes in six animals and with 374-channel Neuropixels probes in two additional animals.

### Single neurons show dynamics at multiple timescales

On fast timescales the spiking dynamics of cortical neurons can be summarised by the interspike interval (ISI) distribution. The characteristic irregular firing of cortical neurons results in ISIs varying by several orders of magnitude (Softky and Koch, 1993), with some neurons also exhibiting ISI histogram peaks indicating rhythmicity at particular frequencies **(Figure 1a).**

**Figure 1.**
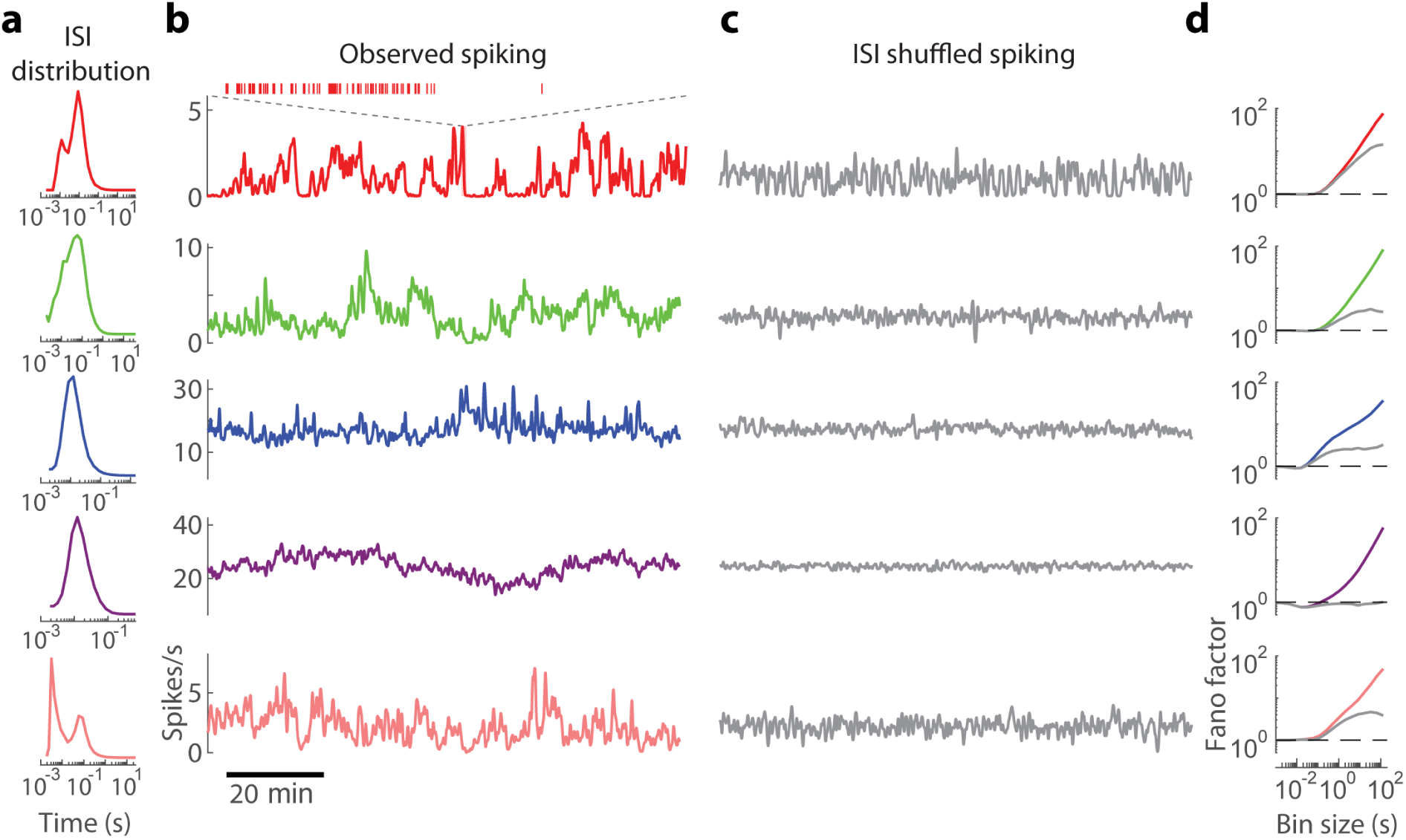
Fast and slow timescale dynamics of individual cortical neurons. **(a)** ISI distribution of five simultaneously recorded example neurons in mPFC of an awake, head-fixed mouse, (b) Firing rate (smoothed with 8 s FWHM Gaussian) over the course of the recording for the neurons in **a. (c)** Firing rate of ISI-shuffled spike trains (cf. b). **(d)** Fano factors of spike counts using bins of 10^−3^ - 10^2^s for original (colour) and ISI-shuffled (grey) spike trains.

A neuron’s ISI distribution was not on its own sufficient to account for the structure of its spike train at long timescales. Indeed, visual inspection shows that cortical firing rates typically fluctuate on timescales of minutes or more **(Figure 1 b),** longer than almost all ISIs of neurons with firing rates ≥ 1 spike/s. Synthetic spike trains created by randomly reshuffling the original ISIs did not have this slow timescale dynamics **(Figure 1c).** The discrepancy between actual and ISI-shuffled data could be summarised by the spike count Fano factor: the variance divided by the mean of spike counts in bins of prescribed temporal duration **(Figure 1d).** For bins of short duration (1-100 ms), Fano factors were close to 1, and the Fano factors of the original and shuffled data were similar. However for bins of 1 −100 s, the Fano factors of actual data were several-fold higher. Across all analysed neurons (n = 775), the Fano factor of the number of spikes in 1024 ms bins was 1.6 times higher in the actual data compared to ISI-shuffled trains, and with 16384 ms bins it was 4.8 times higher (for a summary across all bin sizes see the error of **ISI** model in **Figure 2e,** below). These results demonstrate the major contribution of slow dynamics to spiking variability in the cortex.

**Figure 2.**
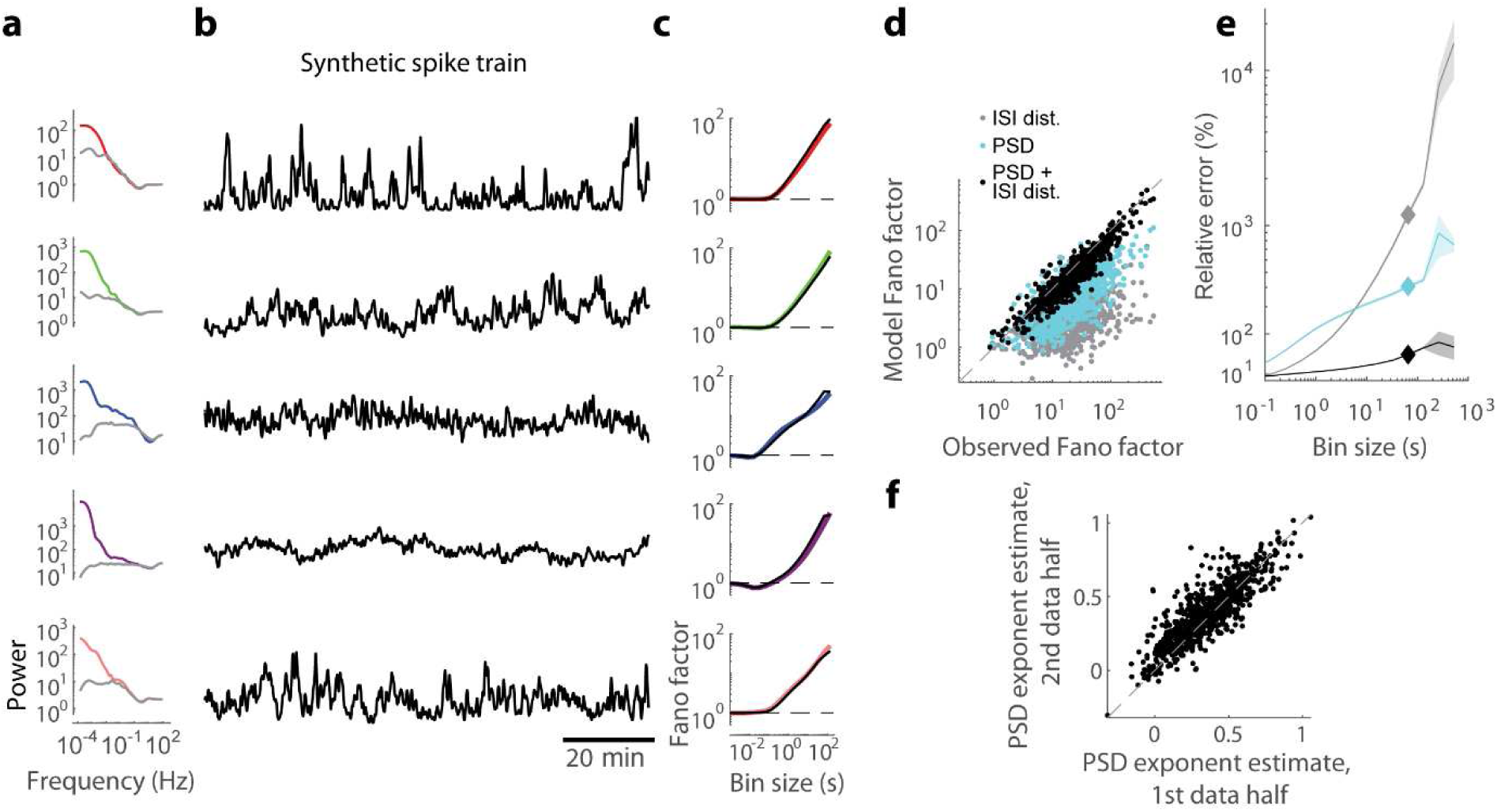
Modelling spiking dynamics on fast and slow timescales. **(a)** PSD of the original (colour) and ISI-shuffled (grey) spike trains for the five example neurons shown in Figure 1. **(b)** Firing rate of synthetic spike trains constructed by requiring that their ISI distribution and PSD match the original data (cf. Figure 1b-c). **(c)** Fano factors of spike counts using bins of 10^−3^ - 10^2^s: the plots for original data (colour) and for synthetic spike trains (black) closely match (cf. Figure Id), **(d)** Observed and predicted spike count Fano factors for 65 s bins for the entire dataset (775 neurons). Predictions were based on ISI distribution only (grey), on PSD only (cyan), or on the full model in which both constraints apply (black), **(e)** Relative error (in %) of predicting the observed Fano factors for bins of 10-^3^ - 10^2^s for the three models, averaged over all neurons. Diamonds mark values for 65 s bin, shown in d. Shaded area shows the standard error, **(f)** The PSD of each neuron was fit with a const/*f*^β^function in the range 0.01-1 Hz. The power-law exponent β is specific to each neuron, which is demonstrated by the fact that the values estimated separately in two halves of the recording closely match (R^2^ = 0.69, P < 1?–^100^).

Although the ISI distribution could not alone capture the infraslow portion of a cell’s spiking dynamics, the combination of ISI distribution and spike train power spectral density (PSD) together provided a good approximation. Because our recordings lasted several hours, we were able to compute power spectra down to very low frequencies, where they typically showed a large peak, indicative of infraslow dynamics **(Figure 2a).** We devised an algorithm that generates synthetic spike trains with pre-specified PSD and ISI distributions (see Methods). The slow-timescale firing dynamics of these synthetic spike trains was visually similar to the original data (compare **Figure 2b** with **Figure 1b)** and closely matched the observed Fano factors over multiple timescales **(Figure 2c),** as expected from the analytical relationship between Fano factor and autocorrelation of a stationary spike train (Teich et al., 1997). The full model was better than models that used either PSD or ISI distribution independently **(Figure 2d).** At slow timescales (e.g. 1 minute; **Figure 2e),** Fano factors predicted from PSD alone are significantly closer to the actual values than ISI-based predictions, but still not as accurate as the full model. For fast timescales (e.g. 100 ms), ISIs predict spike count accurately, but the PSD alone is insufficient **(Figure 2e).** The full model respects both constraints, and as a result provides predictions that are significantly better than either ISI distribution or PSD alone **(Figure 2d-e).**

Cortical neurons are diverse in their intrinsic dynamics. This diversity is well characterised at short timescales by differences in spike regularity (Maimon and Assad, 2009) and by the differing propensity of neurons to emit complex-spike bursts (McCormick et al., 1985; de Kock and Sakmann, 2008), but dynamical diversity at slow timescales is largely unexplored. To address this question, we observed that the PSD of most neurons had power-law profile over the 0.01 - 1 Hz range, with the exponent significantly different between neurons **(Figure S1a).** Fitting spike train power with a const/f**\*^3^** function over 0.01-1 Hz revealed that the power-law exponent β covered a range of 0.39±0.19 (mean and standard deviation for n = 775 neurons), and was conserved when fit separately for the first and second halves of each recording **(Figure 2f;** R^2^ = 0.69 overall; median R^2^ of individual recordings = 0.63; P < 0.05 in each recording). The power law exponent was unrelated to mean firing rate and weakly related to bursting **(Figure S1b-d).** We conclude that cortical neurons are diverse in the strength of their infraslow firing rate fluctuations, and that the structure of these fluctuations can be summarised, to first approximation, by the PSD slope *β*.

### Population coupling strength is unrelated at fast and slow timescales

To understand how the slow dynamics of individual neurons was related to that of the entire population, we started by considering how individual neurons are related to the population rate - the summed rate of all spikes detected on the probe. At short timescales, neurons vary continuously in the strength of their coupling with population rate (Okun et al., 2015). To characterise the relation of neurons to the population across multiple timescales, we extended this analysis into the frequency domain. Analysis in frequency domain does not suffer from the inherent ambiguity of time domain analysis, where correlations computed using a time bin of a specific duration reflect co-modulation not just on the scale of the bin, but also on all slower timescales (Brody, 1999).

The PSD of population rate had 1/f profile, similar to the profile of PSDs of single neuron spike trains. However, in all frequencies < 1 Hz, the population rate PSD was several fold higher than the sum of PSDs of all the individual spike trains that together comprise the population rate **(Figure 3a, Figure S2).** As the PSDs of independent neurons would add linearly, this result indicates that infraslow fluctuations in firing rates of neurons were correlated in all these frequencies. To estimate the coherence between population rate and spike trains of specific neurons, we considered the former as a continuous function of time, and computed its ‘rate adjusted coherence’ (Aoi et al., 2015) with the spike train of each neuron, which accounts for differences in mean firing rate between neurons **(Figure S3a-b;** see Methods). To verify that this method provided a reliable measure, we estimated coherence separately on two halves of single recordings, obtaining similar estimates for coherence on both slow and fast timescales for most neurons (for 0.1 and 10 Hz, correspondingly: overall **R^2^** = 0.63 and 0.75, median **R^2^** of individual recordings = 0.59 and 0.61, P < 0.05 in 19/26 and 22/26 recordings; **Figure 3b-c**, see also **Figure S6).**

**Figure 3.**
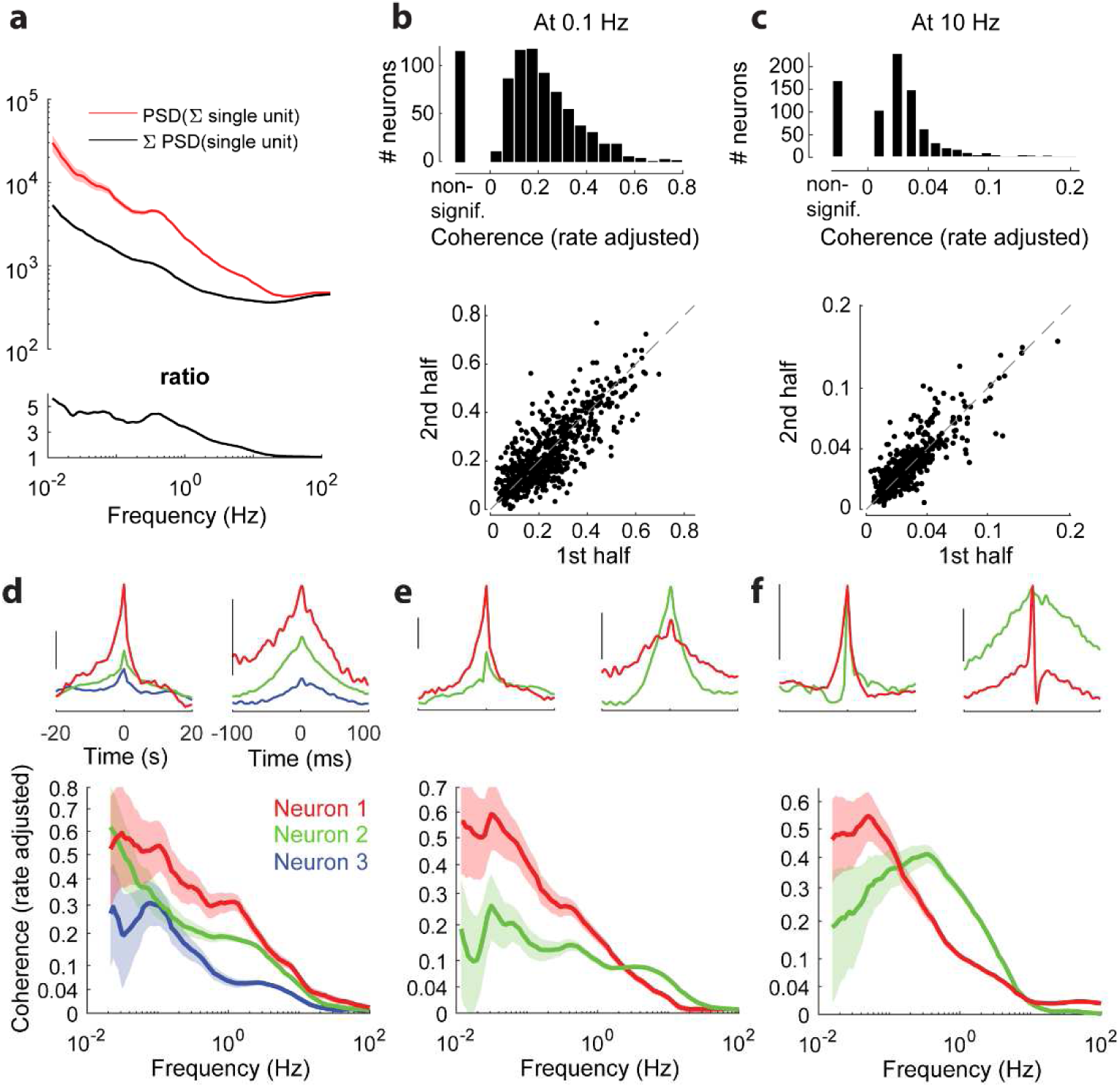
Frequency-resolved population coupling. **(a)** Top: PSD of population rate and sum of PSDs of individual spike trains comprising the population rate, in an example recording. Bottom: the ratio between the two, indicating that the former is several-fold higher, **(b)** Top: Distribution of rate adjusted coherence with population rate at an example slow timescale frequency (0.1 Hz) across all neurons. Some neurons have no significant coherence (“non-signif.”). Bottom: rate adjusted coherence with population rate at 0.1 Hz, evaluated separately in first and second halves of the recordings, R^2^ = 0.63 (P < 10^−100^)**. (c)** Same format as **b,** for fast time scale example frequency (10 Hz), R^2^ = 0.75 (P < 1O^−100^). (d) Top: Time domain correlation between spike trains of three example simultaneously recorded neurons and the population rate, on slow and fast time scales (scale bar: median amplitude of the correlation across all neurons in the recording). Bottom: Rate adjusted coherence of each example neuron with population rate, (e) Two example simultaneously recorded neurons, where one (red) has high coherence with the population in low frequencies and low coherence in high frequencies, whereas the other neuron (green) exhibits the opposite behaviour. Layout as in **d. (f)** Two example simultaneously recorded neurons whose relative strength of population coupling switches twice over the frequency range, furthermore one of the neurons (green) has a non-monotonic coherence with population. Note that the time domain correlation with population rate of both neurons is of equal magnitude on both fast and slow timescales. Layout as in **d.** In **c-f** coherence values are shown using power function scaling, to make low values visible. In **d-f** shaded areas indicate 95% confidence intervals.

Coherence analysis revealed widely diverse relationships to population rate, both between neurons, and between timescales within individual neurons. In all cases, coherence decayed to zero with increasing frequency, owing to the point processes nature of neuronal spike trains **(Figure S3 c-d;** see Methods). However the manner of this decay varied greatly between neurons **(Figure 3 d-f).** In some cases the relative strength of different neurons’ population coupling was conserved across frequencies (e.g. the red neuron in **Figure 3 d** has consistently larger coherence than the green or blue neurons). However, simultaneously recorded neurons often showed different rates of coherence decay: in 25% of simultaneously recorded pairs each neuron had a significantly stronger coherence in a subset of frequencies **(Figure 3 e-f).** Furthermore, some neurons’ coherence with population rate was nonmonotonic (15% of cells, e.g. **Figure 3 f**). On average across neurons, rate adjusted coherence with population rate remained < 0.5 in all our recordings, even in frequencies as low as 0.1 - 0.01 Hz **(Figure 3 b, Figure S4).**

### Phase of population coupling differs across timescales

Coherence is an indication of a constant phase relationship between two processes. Thus, if a neuron has high coherence with population rate at a given frequency, this means it fires at a reliable phase with respect to the population - but does not imply that this phase is zero. Phase analysis showed that most neurons had a stable phase preference with respect to population rate across halves of the recording on both slow and fast timescales **(Figure 4a-b,** see also **Figure** S6). It also revealed a major difference between phases of population coupling on slow and fast timescales, with out of phase activity severalfold more likely in the infraslow range **(Figure 4a-e).** Specifically, at ≥ 10 Hz just 5% of cells had phase closer to ***n*** than to 0, whereas at ≤ 0.3 Hz this was the case for ≥ 28% of the cells (p <10”^28^, Z-test for equality of two proportions). This however did not completely summarise a neuron’s phase preference: even within a mode, there remained significant correlation in a neuron’s precise phase from one half of the recording to the other **(Figure 4a),** and at high frequencies phases also had reliable non-zero values across the two halves of the data set **(Figure 4b,d).** The preferred phase distribution in the infraslow frequencies was not symmetric: more neurons had phases between π/4 and 3π/4 (i.e. leading the population rate) than between **-3 π/4** and -*π /*4 (lagging the population rate, e.g. 16% vs 10% at 0.1 Hz, p < 0.001). The fact that neurons show an asymmetric phase distribution relative to their summed activity might seem contradictory, but was possible because neurons which lagged the population had higher firing rates compared to neurons which led it (4.5 ± 4.8 vs 3.1 ± 4.8 spikes/s, for ***-3n/4*** to - π/4 vs *n/4* to **3π/4,** p < 0.0001). In contrast to the phase preference of individual cells, the relative phase of population rate on different shanks or tetrodes (or on different segments of the Neuropixels probe) was close to 0 in all frequencies **(Figure** S5), suggesting that variations in population coupling phase mainly differ within, rather than between local populations.

**Figure 4.**
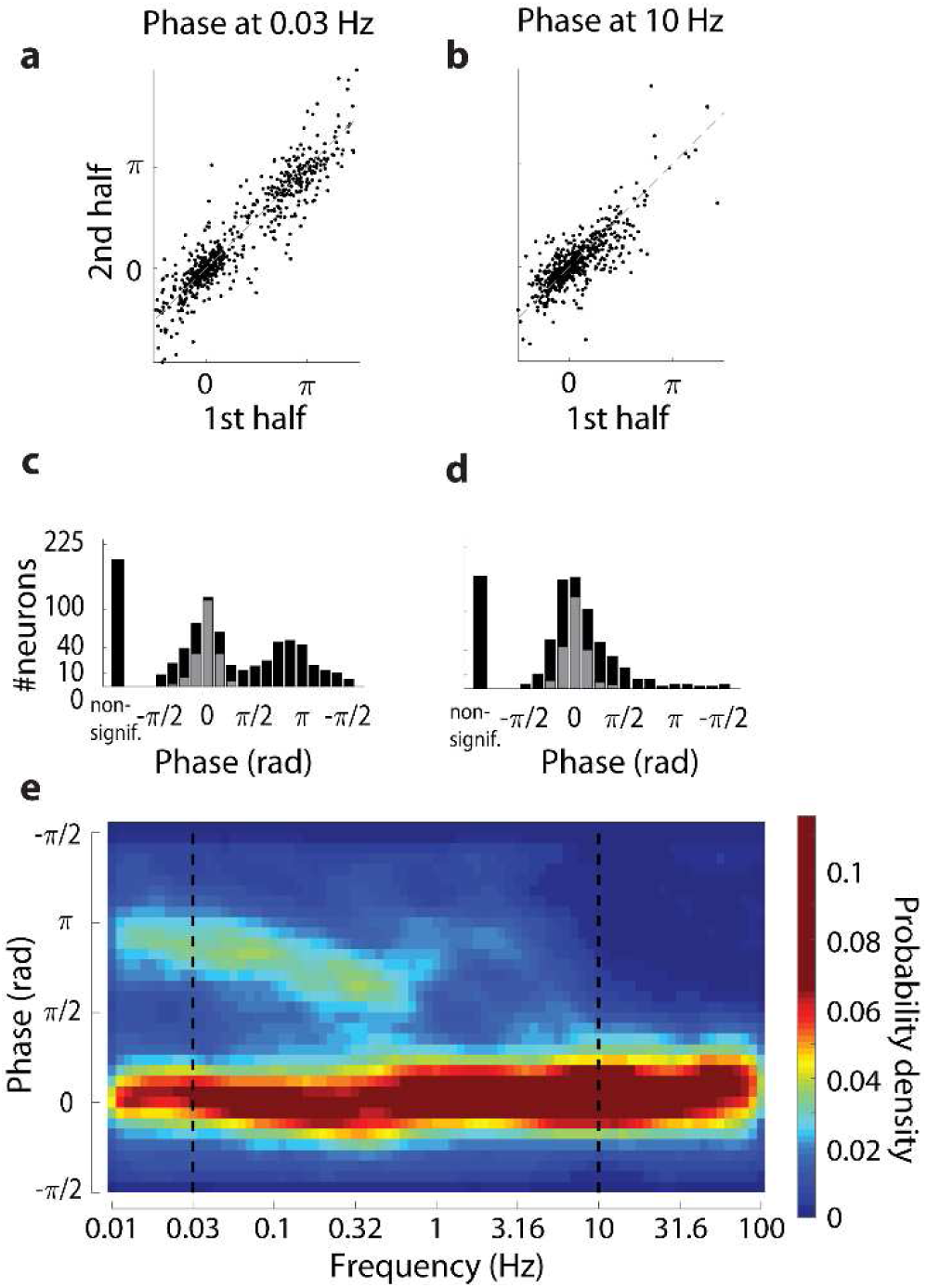
Phase of population-coupling, (a, b) Phase evaluated separately in first and second halves of the recordings, indicating that it is a conserved property for most neurons. Average absolute discrepancy between the two halves: 0.44±0.43 rad and 0.34±0.39 rad (mean and standard deviation for n=582 and n=610 neurons with statistically significant phase preference at 0.03Hz and 10 Hz, correspondingly). Explained circular variance: 0.79 and 0.54 **(P** < 10 ^1S^). **(c, d)** Distribution of the preferred phase of firing of individual neurons with respect to population rate at example frequencies (0.03 and 10 Hz). Some neurons have no significant coherence or phase preference (“non-signif.”). Grey: neurons for which the phase was not significantly (i.e. P > 0.05) different from 0. **(e)** Pseudo-colour histogram of phase preference with respect to population rate across 0.01 - 100 Hz. Dashed lines indicate the two example frequencies shown in **a-d.**

The phase spectra of individual neurons were diverse, and could show a complex dependence on frequency. According to the phase distribution histogram **(Figure 4e)** one expects to find neurons’ phase preference to be close to either 0 or u at infraslow frequencies, and close to 0 at high frequencies. While many neurons conformed to this pattern **(Figure 5a, Figure S6a),** neurons that were anticorrelated with infraslow population rate differed in the dependence of phase preference on frequency: it was discontinuous, with clear subdomains and a drop in coherence in some neurons, but gradual in others **(Figure 5b-c, Figure S6b-c).** We also observed neurons whose phase preference did not fit the overall pattern, e.g., having phase preference of ∼π/2 in infraslow frequencies **(Figure 5d, Figure S6d)** or exhibiting altogether different behaviours **(Figure 5e, Figure S6e).**

**Figure 5.**
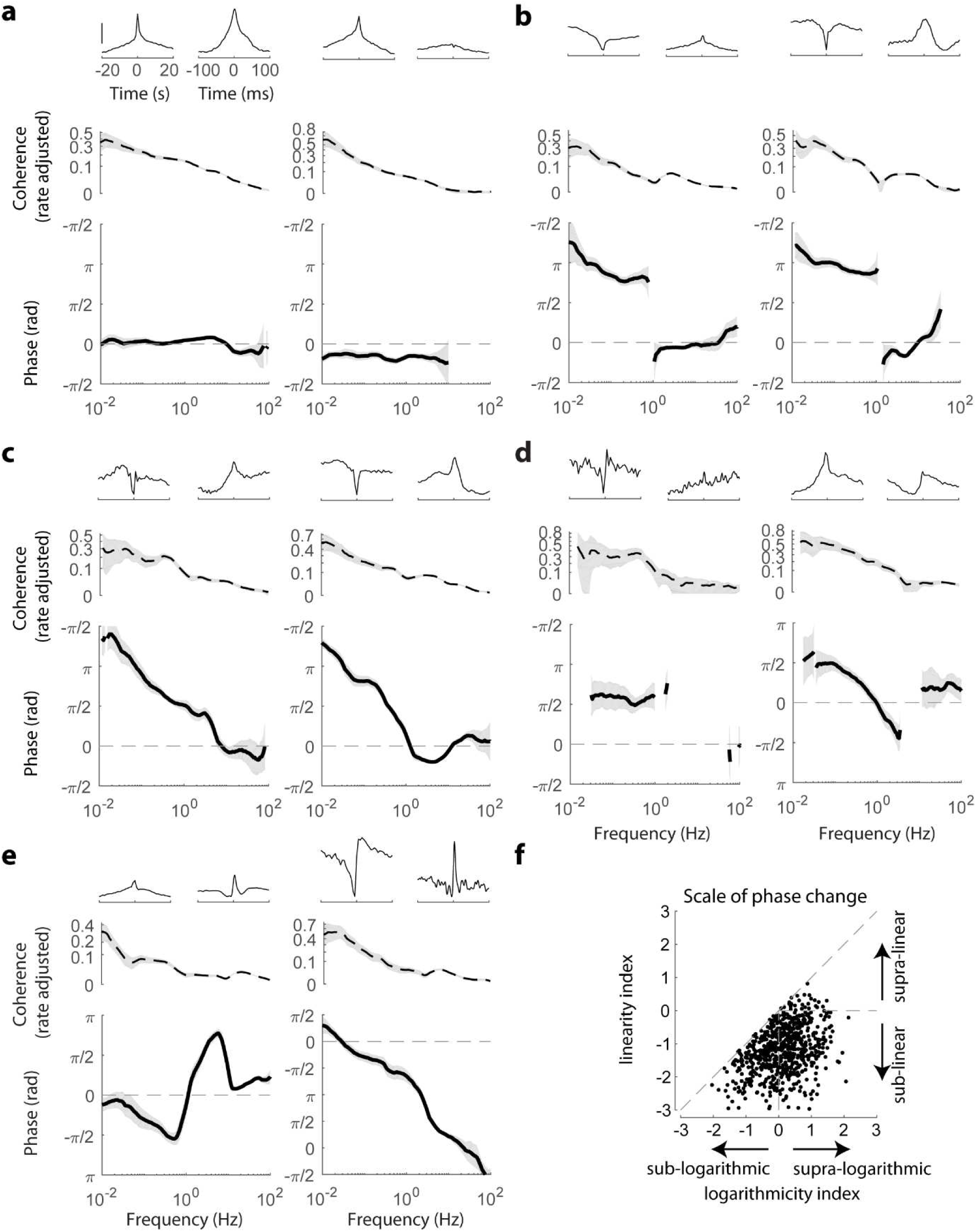
Population coupling phase spectrum. **(a)** Examples of neurons whose firing has phase preference close to 0 with respect to population rate. In the second example the phase is at the same time significantly distinct from 0. Top: time domain correlation between the neuron and population rate on fast and slow timescale (scale bar: median amplitude of the correlation across all neurons in the corresponding recording). Middle: rate adjusted coherence with population rate. Bottom: phase spectrum, (b) Examples of neurons with sharp transition between ∼0 phase preference in high frequencies and ∼π phase in infraslow frequencies, dividing the frequency range into two clear subdomains, **(c)** Examples of neurons whose phase preference is close to 0 in high frequencies and gradually becomes close to *n* in the infraslow frequency range, **(d, e)** Additional examples of observed phase spectra behaviours. Panels **b-e** use the same format as **a,** shaded areas in **a-e** indicate 95% confidence intervals, y-axis of coherence plots uses power function scaling, to make low values visible, **(f)** Logarithmicity index vs linearity index (see Methods) of the longest continuous interval of the phase spectrum of each neuron.

Phase modulation seemed to occur on logarithmic rather than linear scale **(Figure 5c-e).** To assess the rate of phase changes, we devised two indexes which quantify the linearity and logarithmicity of the phases (see Methods). The linearity index is 0 for phase changing on linear scale, and it is positive (negative) for phase changing on supra- (sub-) linear scale. Similarly, the logarithmicity index is 0 for phase changing on logarithmic scale, and it is positive (negative) for phase changing on supra- (sub-) logarithmic scale. We found that phase spectra were overwhelmingly changing sub-linearly (just 4% had positive linearity index), whereas the logarithmicity index values were about equally distributed around 0 (logarithmicity index of 59% of the neurons was positive, **Figure 5f).** We conclude that the phase between single neuron and population rates predominantly changes on a logarithmic scale with frequency.

The logarithmic rate of phase preference change implies that phase in nearby frequencies is similar. In other words, when only a small range of frequencies is considered (e.g. on linear scale), the phase is approximately constant, and thus to a first approximation single cell neuronal dynamics with respect to population rate is scale-free. To test this prediction we compared how well constant phase and linear phase models fit the phase preference in nearby frequencies (0.1 Hz vs 0.03 Hz or 0.32 Hz). The former model corresponds to scale-free dynamics, the latter to a lead or lag by a fixed time interval between an individual neuron and the population rate. As predicted by the logarithmic rate of phase change, the constant phase model fit the data substantially better than the linear model **(Figure** S7).

### Infraslow dynamics correlates with pupil diameter

Head-fixed mice, such as those we recorded here, show fluctuations in alertness levels over time. To address the degree to which the infraslow dynamics we observed could relate to alertness, we monitored the animals’ pupil area in a subset of head-fixed recordings **(Figure 6a).** At 0.03 Hz, 65% (350/541) of the neurons were significantly coherent with the pupil area signal, and the magnitude of this coherence was consistent when estimated from separate halves of the recording **(Figure 6b;** P < 0.01 in 10/13 recordings, the median percentage of variance in one half of the data explained by the other half across recordings; 41%). Phase preferences were similarly stable, and the phase distribution had two clear peaks **∼n** rad apart **(Figure 6c),** consistent with the existence of two populations positively and negatively coupled to arousal (Stringer et al., 2018).

**Figure 6.**
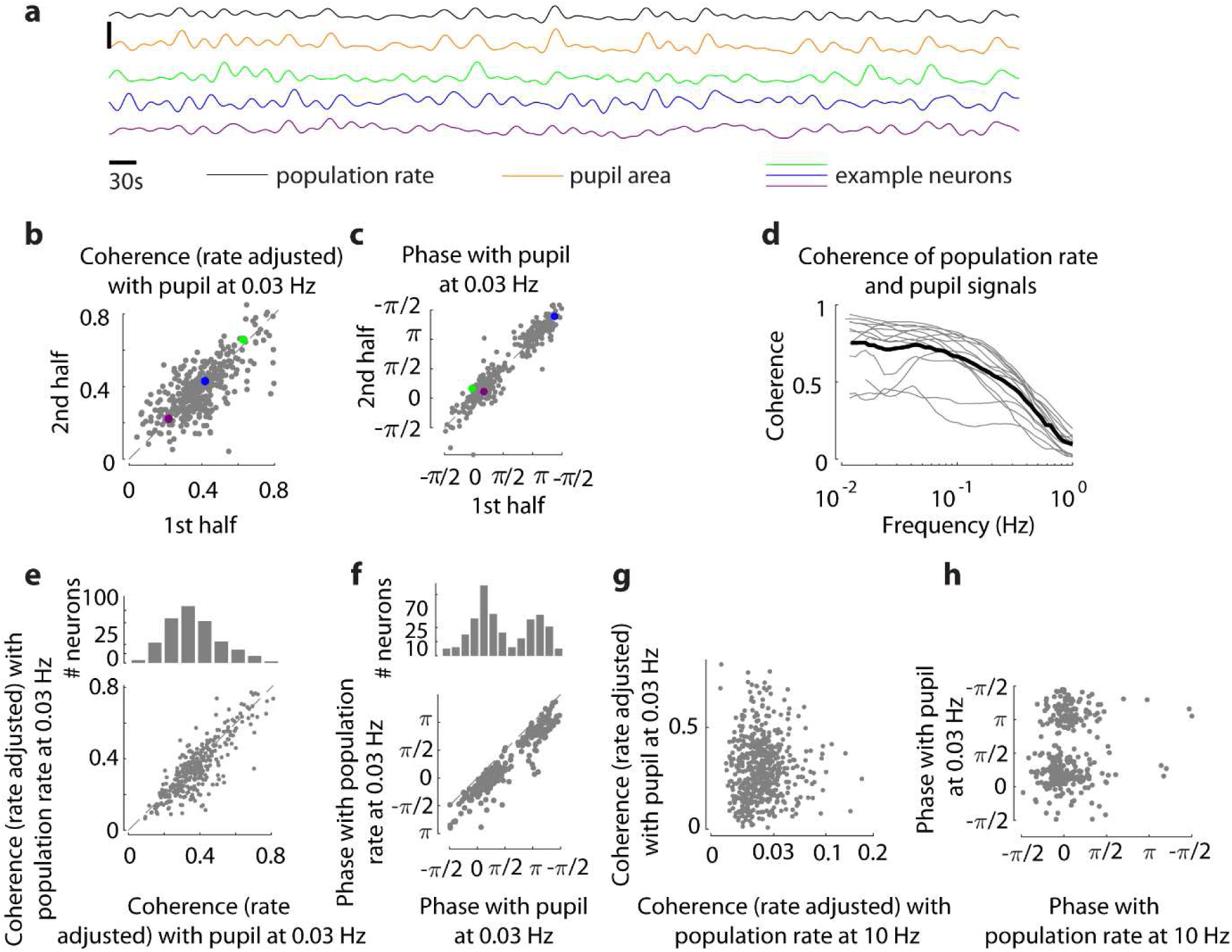
Global origin of infraslow dynamics. **(a)** Population rate, pupil area and spiking activity of example neurons during **1000** s portion of a recording. For presentation purposes only, signals were low-pass filtered below **0.05** Hz and z-scored, vertical scale bar is **5** standard deviations, **{b, c)** Magnitude (rate adjusted) and phase of coherency of individual neurons (n = 350, from **13** recordings) with pupil area signal, separately estimated from two halves of each recording, shown for an example frequency of 0.03 Hz. In **b,** R^2^ = 0.42 (P < 10^−50^). In **c,** average absolute discrepancy between the two halves: 0.34±0.35 rad, 0.88 explained circular variance (P < 10 ^−16^). Coloured dots represent the example neurons shown in **a. (d)** Coherence between population rate and pupil area, in individual recordings (grey, n = 13 recordings in 5 animals) and in their average (black), **(e, f)** Top: Distribution of coherency magnitude (rate adjusted) and phase of individual neurons’ spiking with respect to the pupil area signal at 0.03 Hz. Bottom: magnitude and phase of coherency of individual neurons with pupil area vs their coherency with population rate. R^2^ = 0.67 (P < 10^−90^) in **e,** 0.85 explained circular variance (P < 10^−16^) in **f. (g, h)** Coherence (phase) of individual neurons to pupil signal on slow timescale (0.03 Hz) and their coherence (phase) to population rate on fast timescale (10 Hz) are uncorrelated (P = 0.85, Spearman correlation in **g,** P **=** 0.31, circular correlation in **h).**

Next we considered how individual neurons’ coupling to the pupil and to the local population rate are related. Visual inspection of population rate and the pupil area signals suggested the two are similar in the infraslow range **(Figure 6a),** which was confirmed by coherency analysis showing that the two signals were highly coherent in frequencies < 0.1 Hz **(Figure 6d).** Correspondingly, individual neurons’ coherence with pupil area closely matched their coherence with population rate **(Figure 6e).** Phases with respect to the population rate and the pupil area were also closely matched; a consistent gap between the two (which at 0.03 Hz constituted 0.78±0.51 rad) indicates that neuronal spiking preceded the pupil signal **(Figure 6f).** Importantly, we observed no relationship between coherence with population rate on fast timescales and coherence with the pupil signal (P = 0.85, **Figure 6g;** P > 0.2 in each individual recording, Spearman correlation) and no relationship between the phases (P = 0.31, **Figure 6h,** P > 0.17 in each individual recording, circular correlation; also see **Figure** S8). This observation is consistent with the idea that population coupling on slow timescales is controlled by separate mechanisms from the local synaptic inputs driving fast timescale population coupling.

## Discussion

We used frequency domain analysis to examine the activity of neuronal populations in medial prefrontal cortex (mPFC) across frequencies spanning four orders of magnitude (0.01 - 100 Hz). Our findings point to a fundamental difference between fast and infraslow timescale cortical dynamics. The strength of a neuron’s population coupling at fast and slow timescales was unrelated; furthermore, at fast timescales nearly all neurons fired at preferred phases close to 0 relative to population rate, whereas at slow timescales the phase distribution was bimodal, with preferred phases of ∼30% of the neurons closer to π Population coupling in infraslow, but not fast frequencies reflected coupling to brain-wide arousal signal (pupil area). While these general rules held for most neurons, a great diversity of fine-detailed behaviours was seen within local populations, for example regarding the slow-timescale dynamics as captured by a neuron’s power spectrum, and the way its coherence depended on frequency.

The difference between population coupling at fast and slow timescales likely indicates different mechanisms driving these types of coupling. Fast timescale dynamics reflects local synaptic activity (Haider and McCormick, 2009), and the fast-timescale population coupling of individual neurons is correlated with the number of the local synaptic connections they receive (Okun et al., 2015). In contrast, infraslow dynamics correlates with global, brain-wide phenomena related to arousal, which are controlled at least in part by neuromodulatory inputs (McGinley et al., 2015; Reimer et al., 2016); a similar mechanism has been suggested for the global component of resting state fMRI measurements (Scholvinck et al., 2010; Wong et al., 2013; Turchi et al., 2018). The fact that a neuron’s population coupling on fast and slow timescales were uncorrelated therefore suggests that the degree to which a neuron’s firing is controlled by global brain states is unrelated to its local connectivity; for instance, a neuron weakly affected by neuromodulatory tone could still be strongly wired into the local network. The hypothesis that fast and slow population coupling arise through different mechanisms is also supported by observations of neurons whose phase with population rate had discontinuous subdomains in high and low frequencies, and by the fact that only slow-timescale population coupling phases were bimodal. The latter observation is consistent with prior results: while neurons with weak fast-timescale population coupling were previously described, there are very few with negative fast-timescale coupling (Okun et al., 2015). However, neurons that couple negatively as well as positively to arousal have been reported at least in visual cortex (Vinck et al., 2015; Stringer et al., 2018).

Most of our present day knowledge on infraslow cortical dynamics comes from fMRI studies of resting state activity (Buckner et al., 2013; Raichle, 2015; Foster et al., 2016). Because fMRI provides a blood-oxygen-level dependent (BOLD) signal rather than a direct measure of neural activity, it is limited to measurements on infraslow timescales. Multiple studies have shown that the BOLD signal correlates with population rate (Logothetis et al., 2001; Ma et al., 2016; Mateo et al., 2017), although disagreements on the BOLD signal’s interpretation remain (e.g. see Winder et al., 2017). Our study provides an account of how individual neurons’ activities combine to produce infraslow fluctuations in population rate, and hence in BOLD (to the extent the two are correlated). The low-frequency power of the population rate was 2-5 times larger than it would be if cells were independent of each other **(Figure 3 a, Figure S2**). Because the recorded populations were spread over hundreds of micrometres, this increase would likely have been even higher if the recorded populations were concentrated in a smaller volume. The contribution of single neuron activity to the mesoscale signal is limited in two ways. First, for majority of neurons their coherence with population rate remained relatively low (typically between 0.2 and 0.4) even in the 0.01 - 0.1 Hz range of frequencies **(Figure S4)**, and for some neurons coherence in this range was found to be even lower than for higher frequencies (e.g. **Figure 3f)**. Second, the infraslow fluctuations in firing rate of many neurons were partially or completely out of phase with the population **(Figure 4**). A potential caveat with linking the present work to resting state fMRI studies is the degree to which the activity we observed is in the pure resting state regime (Logothetis et al., 2009; Winder et al., 2017). While it is possible to find short intervals during which a mouse does not move, this is not the case for intervals longer than a few seconds (e.g., typically mice move their eyes every 5-10 seconds). Thus our results should not be viewed as describing pure spontaneous activity (an ideal which is impossible to achieve in practice for infraslow timescales in awake subjects), but pertain to the actual cortical dynamics which is partially driven by intrinsic behaviours.

At the single neuron level, power spectral analysis was consistent with scale-free dynamics in the infraslow frequency range **(Figure 2a, Figure S1).** Such dynamics is typical of neuronal activity on various spatial scales, from fMRI measurements to ion channels, and elsewhere in biology, e.g., organisation of heart beats (Bassingthwaighte et al., 1994). Similar spectra have been reported for retinal and thalamic cells recorded in anaesthetised cats (Teich et al., 1997), and more recently in resting humans (Nir et al., 2008), with a mean power-law exponent of 0.45, close to the 0.39 value observed here **(Figure 2f, Figure S1b).** For an intuitive interpretation of this value consider that for a signal with power spectrum proportional to 1/f^0.4^, 66% of the total power slower than any chosen frequency ω is concentrated in frequencies ≤ **ω/2.** As a result of these slow changes in firing rate of individual neurons, their spike count variance in minute bins was on average ∼10-fold higher than what fast-timescale spiking dynamics alone (i.e. the ISI model) would predict **(Figure 2e).** The interpretation and causes of such scale-free dynamics are controversial. One suggestion is that scale-free behaviour could be caused by single-cell intrinsic mechanisms such as firing rate adaptation (Marom, 2010; Xu and Barak, 2017), which can result in a frequency-independent lead of ~0.2 rad of the output spiking over sinusoidal input currents with periods < 1 Hz (Lundström et al., 2008; Pozzorini et al., 2013). These effects can build up across more complex networks: for example, when rat whiskers were stimulated by white noise on top of which sinusoidal modulation with 0.3 - 0.03 Hz frequency was added, barrel cortex neurons preceded the sinusoidal stimulus envelope by ∼0.8 rad on average, while thalamic neurons were leading by less than half as much (Lundström et al., 2010). An alternative, recently proposed possibility is that infraslow firing rate fluctuations are driven by slow changes in ion concentrations (Krishnan et al., 2018), whereas our data suggests that brain-wide neuromodulatory inputs have a major role in this phenomenon. While the contribution of each of these mechanisms remains to be elucidated, it is likely that their effect on cortical dynamics is particularly complex on intermediate timescales (~1 Hz) where they interact with fast-timescale local synaptic activity.

## Materials and methods

### Electrophysiological recordings

All experimental procedures were conducted according to the UK Animals (Scientific Procedures) Act 1986 (Amendment Regulations 2012). Experiments were performed at University College London (UCL) under personal and project licenses released by the Home Office following institutional ethics review. Adult C57BL/6 mice of both sexes were used.

The experimental procedures for chronically implanting Neuronexus and Neuropixels probes were previously described in (Okun et al., 2016; Jun et al., 2017). Briefly, in an initial surgery under isoflurane anaesthesia animals were implanted with a custom built head-plate. Following full recovery and acclimatisation to head-fixation, probe implantation was performed under isoflurane anaesthesia. The probes were implanted through a craniectomy above medial prefrontal cortex (0.5 lateral and 1.8 anterior to bregma). Neuronexus probes (A2−2-tet with CM16LP connector package and Buzsaki32 with CM32 connector package) were lowered 1.7mm, placing the recording sites in the prelimbic cortex (PrL). Neuropixels probes were lowered ∼3.5mm (so that the most superficial of the 374 recording sites remained outside of the brain, while the deepest sites were ∼3.5mm inside the brain; the recording sites were thus placed in the cingulate, prelimbic, and infralimbic cortices). The probes were oriented approximately parallel to the cortical layers, 0.5mm lateral offset of the insertion point relative to the midline implied that the probes resided in cortical layers 5 and 6 (which was also confirmed histologically).

Recordings were performed over the course of several months following the probe implantation. For head-fixed recording, mice were placed inside a plastic tube where they could comfortably sit or stand. The recordings lasted 1.5-3 h. In animals implanted with Neuronexus probes, recordings were performed using OpenEphys (www.open-ephvs.org) recording system (Siegle et al., 2017). Mice with a Neuropixels probe were recorded using SpikeGLX system (github.com/billkarsh/SpikeGLX) developed at .ianelia Farm. (Some of the mice were trained and recorded in a behavioural task which the animals would perform for water reward; the data analysed here is from recordings of ongoing activity in separate sessions without behaviour, performed on days when the animals were not water deprived).

Recordings in freely behaving animals implanted with Neuronexus probes lasted 4-8 h. Mice were briefly head-fixed to allow attaching the amplifier head-stage to the probe and then released into their home cage, where they were free to engage in any activity of their choice, while being monitored to make sure that the thin cable leading from the amplifier to the OpenEphys box was not entangled.

### Spike sorting and drift contamination

Spike sorting of Neuropixels recordings was performed using Kilosort software (Paclitariu et al., 2016), with manual curation performed using phy (github.com/cortex-lab/KiloSort and github.com/kwikteam/phy). Spike sorting of Neuronexus probe recordings was performed similarly, or using SpikeDetekt, KlustaKwik and Klustaviewa software suite (Rossant et al., 2016).

We have evaluated the quality of spike sorted units using isolation distance metric (Schmitzer-Torbert et al., 2005) and by quantifying the contamination of the refractory periods of the spike autocorrelograms, which was expressed as proportion of the number of spikes in the first 2 ms of the autocorrelogram relative to the autocorrelogram asymptote (Harris et al., 2000). We have limited the analysis to units with isolation distance > 20 and refractory period contamination < 0.2. Our analyses yielded quantitatively similar results when more (and less) stringent criteria were applied.

A possible concern is that our results, instead of reflecting the properties of actual infraslow fluctuations in the firing rates of cortical neuronal populations, are dominated by contamination introduced by unstable recordings. Such concern is not unique to the present work, and was raised in the past regarding estimation of pairwise correlations, e.g., (Ecker et al., 2010). Here, Neuropixels recordings provided an unprecedented opportunity to detect and monitor drifts, as the recording sites span a contiguous stretch of > 3mm. For spikes detected simultaneously on several contacts, we have computed the vertical location of the ‘centre of mass’ of the spike, according to the relative amplitude of the spike waveform on each contact. Changes in these locations over time, particularly for high-amplitude spikes, reveal potential drifts. In the example shown in **Figure S9,** multiple drift events are visually apparent. In each event, the vertical location of high-amplitude spikes at one particular neighbourhood of the probe drift ∼10 (μ,m upwards over the course of ∼5 s, and over the next ∼20-40 s return to their original location. Similar drift pattern occurs ∼200 μm further down the shank **(Figure S9b),** which is a strong indication that these two drifts are produced by a vertical movement of the probe with respect to cortical tissue, rather than any other cause. In fact, drifts were simultaneously observed at > 10 locations across the top 800 μm of the probe. Drifts were not observed when vertical location of low-amplitude spikes was considered. This is consistent with the idea that a vertical movement has a much larger impact on the waveforms of high-amplitude spikes originating in neurons abutting the probe compared to the low-amplitude spike waveforms of neurons that are more (horizontally) distant.

In the example recording, drifts of high-amplitude spikes were not observed on contacts deeper than ∼800 μm **(Figure S9b).** In recordings from this mouse, all data originating from the top 1 mm was excluded from the analyses. No similar drifts were observed in Neuropixels recordings from the second animal. In recordings performed with Neuronexus probes, the recording sites were located only at the bottom 200 μm of probes which were lowered 1.7 mm into the brain, thus to the extent that drifts of Neuropixels and Neuronexus probes are similar, we do not expect to find vertical drifts in these recordings (since the recording sites were not covering a contiguous interval, the above drift analysis cannot be repeated for Neuronexus recordings).

An additional observation suggesting that our results are not driven by drifts concerns the relationship between amplitude of the different units and their phase preference. If drifts introduce a strong bias into our estimation of phase with respect to population rate, then there might exist some consistent relationship between the phase and spike waveform amplitude of the different units, because drift bias is expected to be stronger for units close to the probe and having high-amplitude spike waveforms. However, no significant correlation between phase and spike amplitude was found in our data. We conclude that drift is an important caveat that has the potential to bias measurements of spiking activity obtained with extracellular probes, however in view of the control analyses explained above, we believe that the phenomena described here are not due to such drifts.

### Pupil tracking

Pupil area was tracked as previously described in (Burgess et al., 2017). Briefly, a camera (DMK 21BU04.H or DMK 23U618, The Imaging Source) with a zoom lens (ThorLabs MVL7000) was focused on one of the eyes of the animal. The eye was illuminated by an infrared LED (SLS-0208A, Mightex). Videos of the eye were acquired at ≥ 30 Hz. In each video frame, excluding frames with blinks, an ellipse was fit to the pupil image, and pupil area was estimated based on this fit.

### Single spike train analysis and modelling

Power spectral density (PSD) of individual spike trains **(Figure 2a)** was estimated using mtspectrumsegpb function of Chronux toolbox (Mitra and Bokil, 2007) <u>(chronux.org)</u>. PSD in different frequencies was estimated by breaking the entire recording into segments of appropriate length, and averaging across them. Specifically, segments for estimating PSD in frequency co Hz had length of at least **7/ω** and at most **10/ω** seconds. For frequencies < 0.01 Hz the spectrum of the entire recording was computed using `mtspectrumpb` function without breaking it into segments. For presentation purposes only it was further smoothed using Matlab’s `smooth` function.

The spike train model which captures both the fast and slow timescale dynamics of cortical spiking relies on both ISI distribution and PSD of spike trains **(Figure 2).** The goal of the model is to generate synthetic spike trains satisfying both types of constraints simultaneously. Let 𝔍 denote the observed ISI distribution of a spike train. For modelling, 𝔍 was represented by a histogram with logarithmically spaced bins of all the observed ISIs (32 bins were used to describe ISIs, from 1 ms up to 200 s). Instead of using the PSD of the spike train itself, the model uses the PSD of the underlying continuous firing rate intensity, which we denote by ***𝓅*** (the two are closely related but distinct, as will be explained in more detail later). For modelling, ***𝓅*** was represented by the power of a continuous signal obtained by convolving the observed spike train with a 50 ms FWHM Gaussian (50 parameters were used to represent *𝓅).* With 𝔍 and *𝓅* as its inputs (82 parameters in total), the goal of the model is to generate a synthetic spike train whose ISI distribution and PSD are as close as possible to the original spike train. An intermediate step towards this final goal is constructing a continuous firing rate intensity signal **r(t)** for the synthetic spike train. However, we start by constructing a different firing rate intensity signal, ri(t), by sampling ISIs from 𝔍 and convolving the resulting spike train with a 50ms FWHM Gaussian. Typically *r*_*1*_**(t)** will have much less power in the infraslow frequencies than what is required. Thus, the first constraint which *r*(*t*) must satisfy is to have power **𝓅.** A straightforward way to generate a signal with a given PSD (using inverse Fast Fourier transform) produces a signal whose values are normally distributed with 0 mean, which is inappropriate for a firing rate intensity function. To work our way around this problem, we require the distribution of values of *r*(*t*) to match the distribution of values of *r*_1_ (*t*). We used an iterative algorithm of (Schreiber and Schmitz, 1996) to generate *r*(*t*) given the constraints on its power and distribution of values. Once *r*(*t*) was generated, we sample a spike train *n*_1_ using *r*(*t*) as a time dependent firing intensity signal. In the final step the ISIs of the spike train are adjusted to have the desired distribution 𝔍. Specifically, we convert the sequence of ISIs in into a sequence of ISI ranks, by replacing each ISI with its rank among all the ISIs of *n*_*1*_. We build the final output spike train n by sampling from 3 the same number of ISIs found in n_1_, and rearranging them according to the sequence of ranks from *n***_1(_** i.e., the ISI rank sequences of *n* and of % match.

When the model is used to generate an output without an explicit constraint on 3(i.e. only ***𝓅*** input is provided), it implicitly assumes 𝔍 has an exponential distribution, with an additional constraint forbidding ISIs < 2ms (representing a hard refractory period).

### Time domain population coupling on fast and slow timescales

Time domain correlation between spike trains of single neurons and the population rate **(Figure 3 b-d)** was computed as previously described in (Okun et al., 2015). Specifically, we computed the inner product between the vectors representing the population rate and single unit spike train at different lags (using Matlab’s `xcorr`), and normalised it by the number of spikes of the single neuron. For fast timescale correlation, the vectors were at 1 ms resolution, and the single neuron spike train was smoothed with Gaussian of 12 ms halfwidth. For slow timescale correlation, the vectors were at 1 s resolution. In both cases the baseline (average values 800 - 1000 ms away from zero lag for fast timescale correlation, and average values 12 - 20 s away from zero lag for slow timescale correlation) were subtracted.

### Coherence analysis

For analysing the relationship between spike trains of individual units and population rate, the latter was obtained by summing all the spikes detected on all the shanks/tetrodes barring the one on which the single unit was recorded. For Neuropixels recordings, where the entire probe consists of one shank (with 374 recording sites over ∼3.5 mm) this approach was not applicable. Instead, for each unit we have performed our analyses with population rate based on all spikes on the probe (except for those of the unit itself) and with population rate based only on spikes from recording sites > 60 μm away from the location of the single unit. All results were almost identical for both conditions. The population rate typically was >100 spikes/s.

Coherence between population rate or pupil area and individual units was estimated, together with its confidence interval, using `coherencysegpb` function of the Chronux toolbox (estimating coherency using `coherencysegcpb` where population rate was considered a continuous signal rather than spike count produced identical results). As in the case of PSD estimation, coherence in different frequencies was estimated by breaking the entire recording into segments of appropriate length for each frequency.

Unlike the more familiar case of coherence between a pair of continuous processes, coherence between a continuous process and a point process (such as a spike train of a neuron) depends on the PSD and the rate of the latter. This (mathematically unavoidable) fact has two implications. First, the *1****/****f* profile of firing rate PSD implies that coherence of the spike train with population rate falls with frequency, even when coherence between the underlying firing rate intensity and the population rate does not. Second, because coherence depends on the rate of the spike train, two neurons whose firing rate intensities are exactly proportional but unequal do not have the same coherence with population rate. To account for this second issue of rate dependence, we use a correction factor to produce a coherence which reflects a firing rate of 1 spike/s, rather than the actual firing rate of the neuron.

More formally, let *y*(*t*) be a continuous process, *n*(*t*) a point process such as a spike train of a single neuron, and λ(t) the intensity of the spike train, i.e., we assume that n(t) is a doubly-stochastic Poisson process with conditional intensity λ(t). It holds that

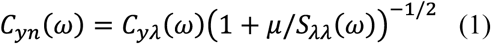

where *C*_*yn*_ and *C*_*yλ*_ denote the coherence between *y*(*t*) and *n*(*t*) or λ(*t*), *μ* is the mean rate of *n*(*t*), and *S*_*λλ*_ is the power spectrum of *λ*(*t*) (Aoi et al" 2015). From the above equation it is clear that two spike trains with proportional but unequal rate intensities (i.e. if *λ*_1_(*t*) **=** *aλ*_*2*_(*t*) where *a* ≠ 1) will have different values for coherence. This is not desirable, therefore instead of reporting the coherence between a spike train and the population rate, we report ‘rate adjusted coherence’ which reflects the coherence that would have been measured if the neuron had a firing rate of 1 spike/s, i.e., if its firing intensity was *λ*(*t*)*/μ* instead of λ(t), see **Figure S3 a-b** for an example. We use a correction factor of (1 + (λ — l)λ/S_nn_(ω)))^−1^/^2^, as derived in (Aoi et al., 2015), to obtain the rate adjusted coherence. The PSD of the spike train, used for the correction was estimated as described above.

The rate adjusted coherence still depends on the PSD of the spike train of the single unit. For instance, it is possible to have two intensity functions λ_1_(t) and λ_2_(t) with equal means of 1 spike/s, and equal coherence with y(t), but with different power spectra. In this case, equation (1) implies that if n_x_ and n_2_ are spike trains sampled according to λ_1_ and λ_2_, then 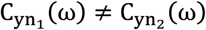 even though 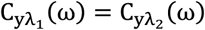. Here, we did not attempt to remove this dependence, which would have required an accurate estimate of Sλλ(ω). In practice, Sxx cannot be directly inferred from S_nn_ because the assumption that n(t) is a doubly-stochastic Poisson process with intensity n(t) does not hold. For example, the existence of refractory period in n(t) reduces the power in all low frequencies in S_nn_(ω) (Bair et al., 1994; Rivlin-Etzion et al., 2006). Furthermore, the spiking of actual neurons is driven by changes in the subthreshold membrane potential V_m_(t) which is rather distinct from λ(t), as exemplified in **Figure** S3 . Of note, this discussion primarily applies to high frequencies, whereas in low frequencies μ is significantly lower than S_λλ_(ω) (or S_nn_(ω)) and thus the discrepancy between C_yn_(ω) and *C*_*yλ*_(ω) is minor (see Equation 1).

To compare the strength of population coupling of two simultaneously recorded neurons across all timescales, we have compared their rate adjusted coherences in the following 9 frequencies: 0.01, 0.03, 0.1, 0.32, 1, 3.2, 10, 32 and 100 Hz. If the null hypothesis that first neuron has higher rate adjusted coherence in these 9 frequencies could be rejected at p ≤ 0.001 (after using Bonferroni correction for performing 9 comparisons), and the reverse null hypothesis could also be rejected with p ≤ 0.001, the two were considered as (a positive) example of a simultaneously recorded pair of neurons where neither neuron dominated the other across all frequencies.

### Phase analysis

Phase of spiking of single units with respect to population rate or pupil area was estimated using the same Chronux toolbox functions used to estimate the coherence (see above). As in the case of PSD estimation, coherence in different frequencies was estimated by breaking the entire recording into segments of appropriate length for each frequency. After the phase in each segment was estimated, circular mean and standard deviation were computed. If the distribution of phases (across the segments) had no statistically significant (at p ≤ 0.05) mean, no phase was assigned (e.g. the non-significant neurons in the histogram in **Figure 4a).**

All phases are specified with respect to the population rate, e.g., a phase of — π/4 means that the single unit lags behind the population rate, whereas a phase of π/6 means that the unit leads it.

### Linearity and logarithmicity indexes

For a continuous, non-constant function *f*(*x*) defined on an interval [*a, b*] (0 < *a < b),* consider the following expression:

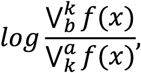

where 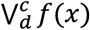 denotes the total variation of *f*(*x*) on the interval [c, *d*]. We define the linearity index of *f(x*) as the value of this expression for *k =* (a *+ b)/2.* Similarly, the logarithmicity index is the expression’s value for 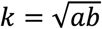. The rationale for these two definitions is that for a function changing on a linear scale, total variation in the first and second halves of [*a,b*] is expected to be of comparable magnitude. Thus, linearity index is close to 0 for functions changing on linear scale (the function itself does not have to be linear, e.g., sin(x) on any sufficiently long interval), positive for supra-linear functions, and negative for sub-linear functions. On the other hand, for a function changing on a logarithmic scale, the total variation in 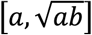 and 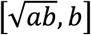 intervals is expected to be of comparable magnitude, thus its logarithmicity index would be close to 0 (while its linearity index would be negative).

For empirically measured *f*(*x*), total variation is contaminated by measurement noise. To avoid this problem, and relying on the fact that phase functions were either monotonic or had just a few extremum points (typical examples shown in **Figure 5),** we used the following expression instead of the one given above:

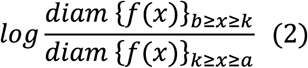

*diam* {*f(x)*}_*d≥x≥c*_ denotes the diameter of the set {*f*(x)}_d≥x≥c_ (except for cases when *f(x)* wraps around *±π,* this is equal to max *f*(*x*) — min *f*(*x*)). For each neuron, expression (2) was evaluated using the longest continuous interval of frequencies over which phase was well-defined (i.e., it had no frequencies in which coherency with population rate was not statistically significant). Neurons for which such interval spanned less than an order of magnitude were excluded from the analysis.

## Acknowledgements

We thank Michael Krumin for the pupil analysis code, and Charu Reddy and Miles Wells for help with histology. M.O. was funded by Springboard award (SBF002Y1045) supported by The Academy of Medical Sciences and the Wellcome Trust, and by BBSRC New Investigator grant (BB/P020607/1). N.A.S was funded by postdoctoral fellowships from the Human Frontier Sciences Program and the Marie Curie Action of the EU (656528). A.L. was funded by Sir Henry Wellcome Fellowship 106101/Z/14/Z. K.D.H. was funded by Wellcome Trust Investigator grants 102264,205093, ERC grant 694401, and Simons Foundation grants 325512, 552341.

## Supplementary figures

**Figure S1.**
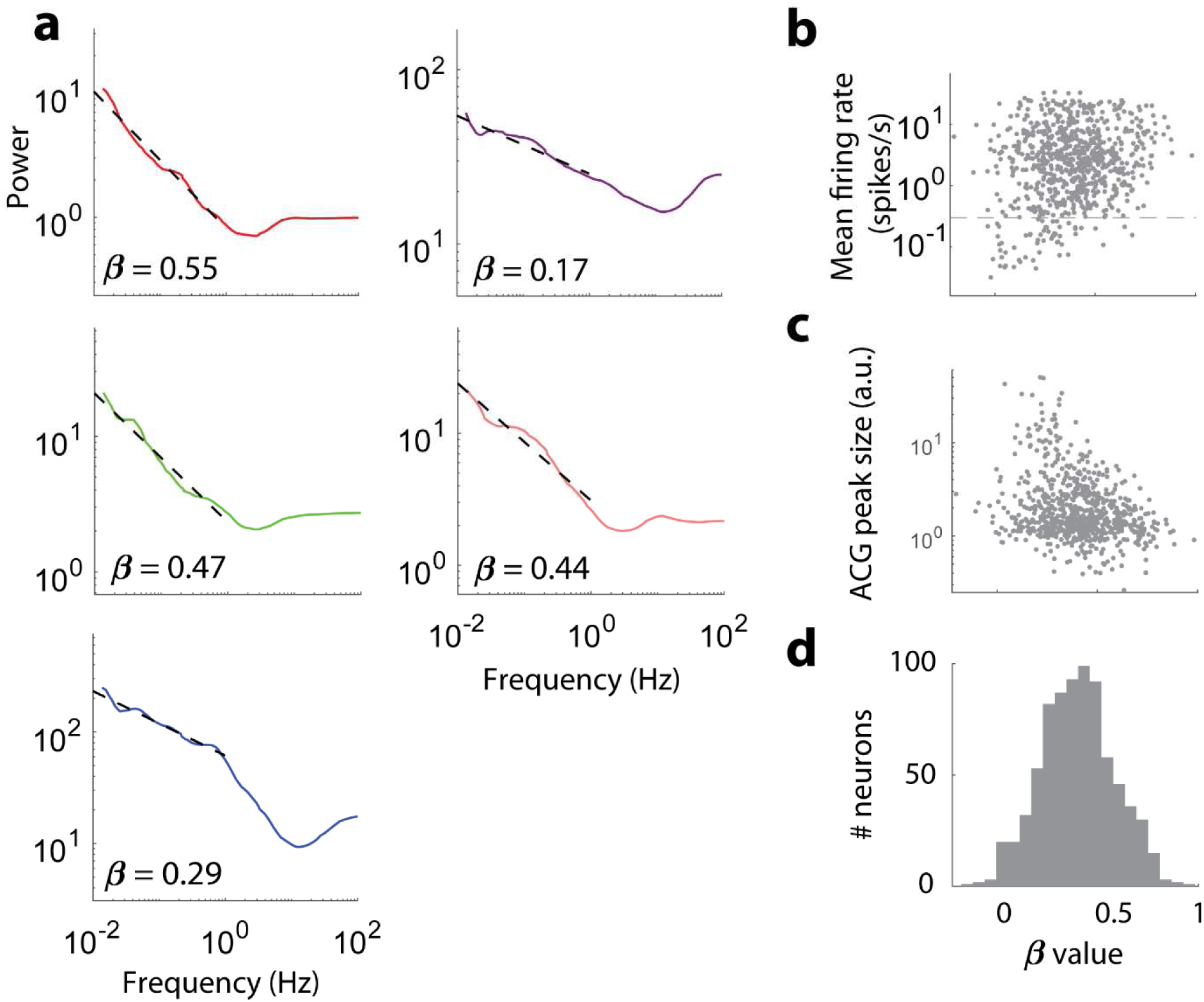
Power-law behaviour of infraslow power spectrum of spiking in cortical neurons. **(a)** The spike train power spectrum in the 0.01 - 1 Hz range was fitted with const/f^*β*^ function. The fit for the five example neurons from Figure 1 is shown by a dashed line, **(b)** Power-law exponent shows no relationship to mean firing rate of the neurons, except for neurons with very low firing rate (where *β* is low owing to estimation bias, equally present in simulated data). For neurons with mean firing rate ≥ 0.3 spikes/s the correlation with *β* was low and insignificant: 0.03, P = 0.39 (Spearman correlation), **(c)** Power-law exponent is weakly correlated with burstiness (the ratio between the peak and baseline of a neuron’s autocorrelogram), r = −0.23, P < 10^−9^ (Spearman correlation). **(d)** Distribution of the power-law exponent value across all the analysed neurons.

**Figure S2.**
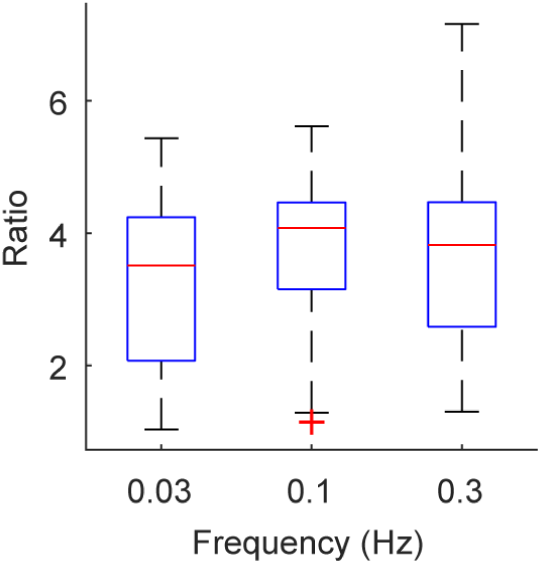
Increase in population rate PSD produced by neuronal coherence. The ratio between PSD of population rate and sum of PSDs of the firing rates of individual neurons that constitute it, averaged across all recordings (n = 26), at 0.03, 0.1 and 0.3 Hz. In these frequencies PSD of population rate was on average 3-4 times higher than what it would have been if the firing rates of the neurons were uncorrelated.

**Figure S3.**
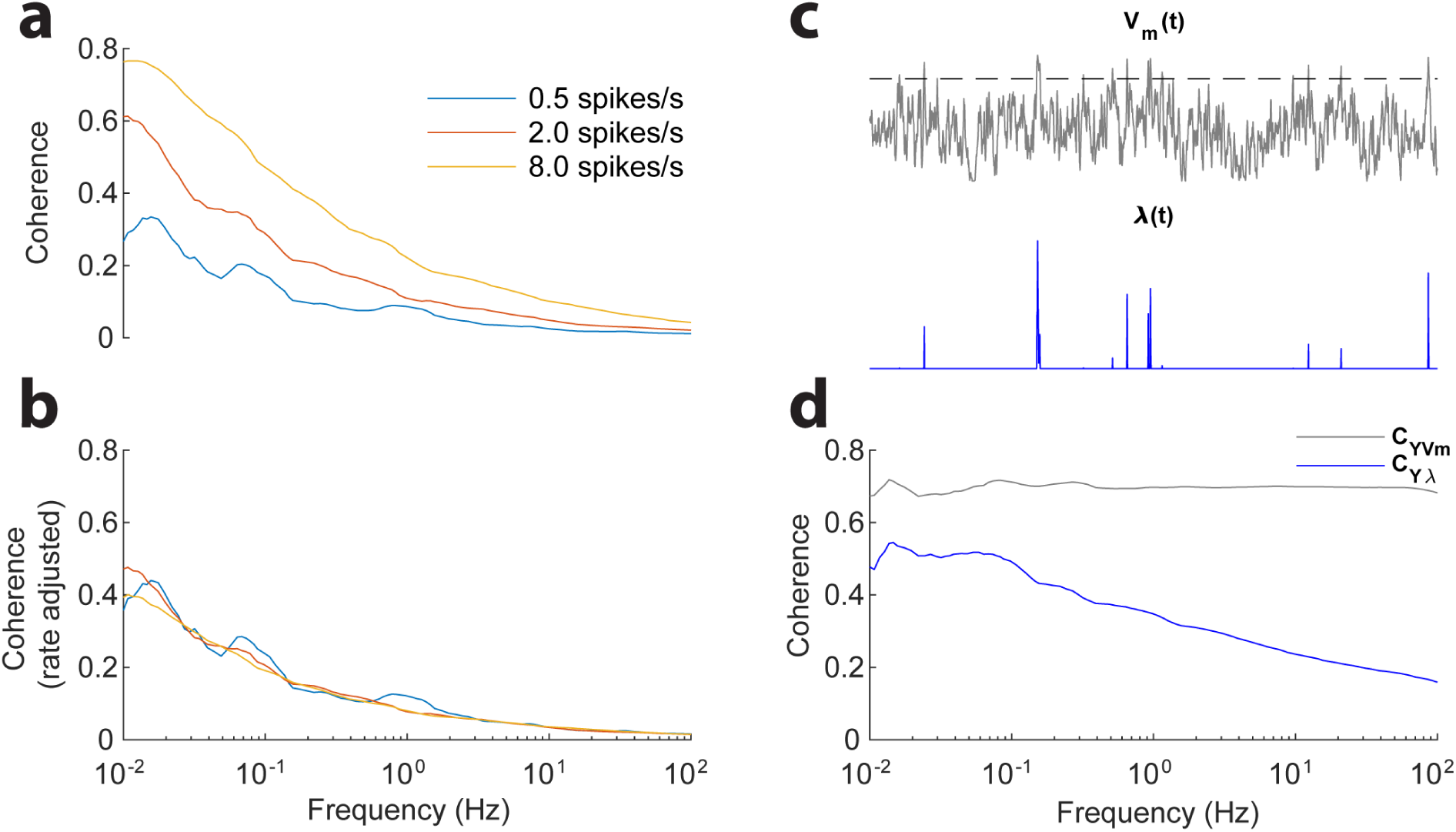
Estimating coherence with spike trains. **a)** Spike trains with rates of 0.5, 2 and 8 spikes/s, and conditional intensity of aA(t), where a controlled firing rate and A(t) > 0 was an artificial signal with 1 /*f* power, were generated. The coherence of the three spike trains with 2(t) depends on their firing rate, although the coherence of their underlying intensity with/l(t) is 1 in all three cases, (b) The rate adjusted coherence of the spike trains in **a** is similar. **(c,d)** Rate adjusted coherence, demonstrated in b, relies on the mathematical formalism of a doubly stochastic Poisson process, which does not apply to actual neurons where spikes are driven by membrane potential (*V*_*m*_) fluctuations, and the spike generation mechanism is to a large extent reliable (Mainen and Sejnowski, 1995). Yet, even for actual neurons one could think of a continuous firing rate A(t) that gives rise to the observed spike train. To a first approximation such A(t) is *V*_*m*_*(t*) transformed through a static non-linearity (determined by the spiking mechanism), as demonstrated by a synthetic example in **c.** Such transformation implies that coherence between any other signal *Y(t)* and A(t) (we will denote it by CVa(A0) deviates markedly from *C*_*YVm*_*(ω)* (the coherence between *Y(t*) and *V*_*m*_(*t*)). In fact, one can show that in this case 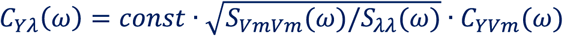, where *S*_*VmVm*_(ω) and *S*_λλ_(<y) denote the PSD of *V*_*m*_*(t*) and A(t). In the example in **c,d,** a pair of artificial signals *Y(t)* and *V*_*m*_*(t)* have a constant coherence of 0.7 across all frequencies, whereas the coherence between *Y(t*) and *λ*(*t*), derived from *V*_*m*_(*t*) via static noniinearity, is no longer constant but falls with frequency, which is explained by the square root of the ratio between PSDs of *V*_*m*_(*t*) and *λ*(*t*).

**Figure S4.**
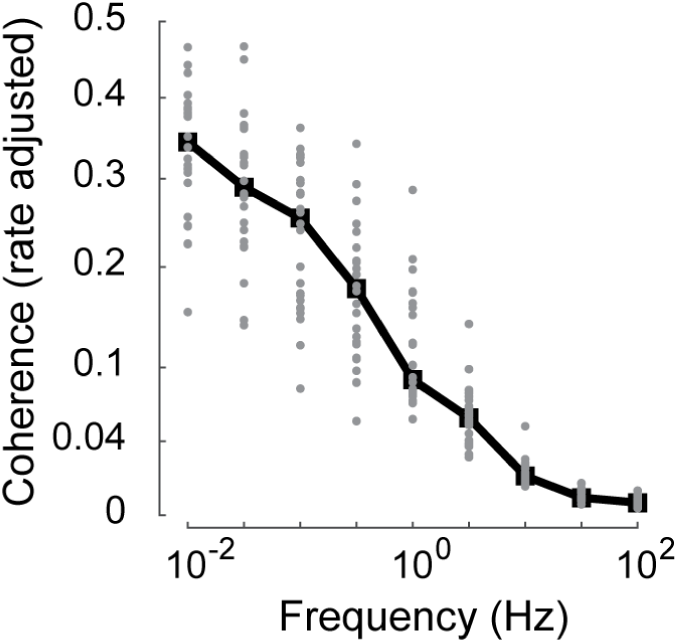
Average frequency-resolved population coupling. The value of rate adjusted coherence with population rate averaged across all neurons in each recording (grey points), and its median across all recordings (black).

**Figure S5.**
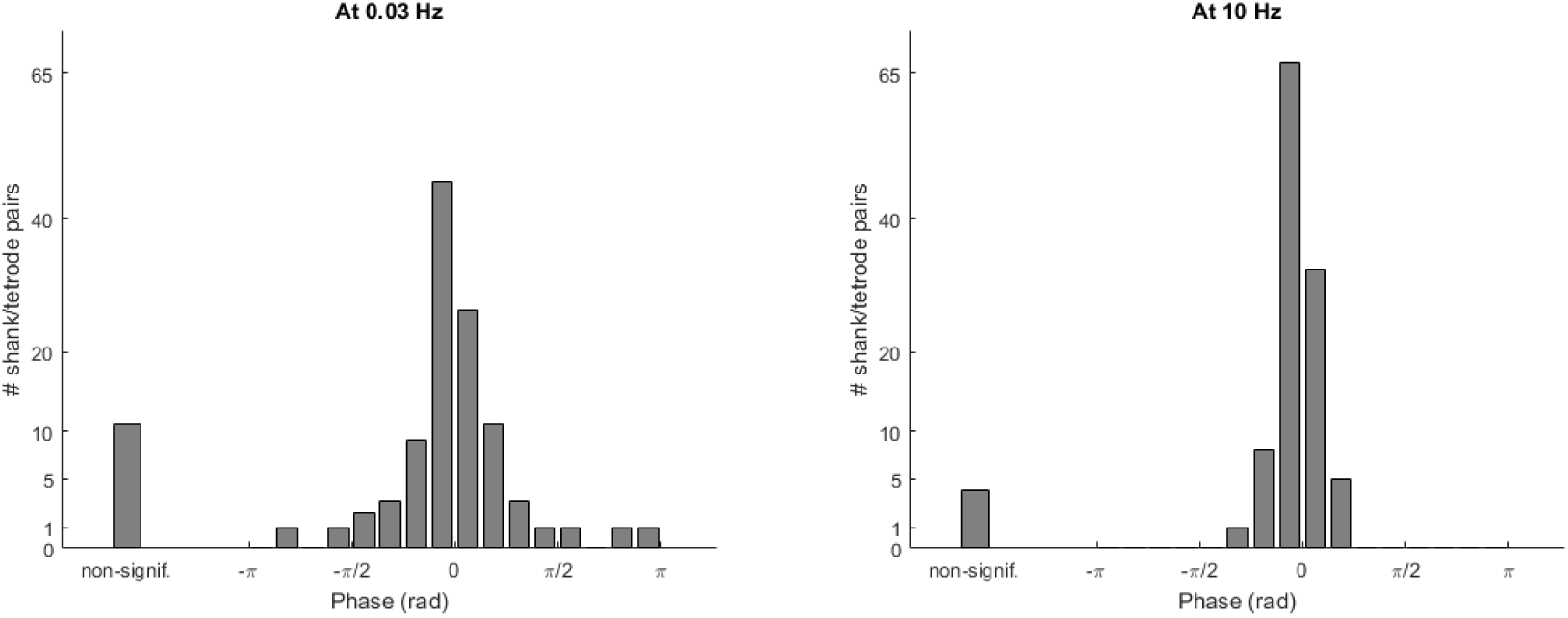
Phase between multiunit activity (MUA) on different shanks or tetrodes in the same recording. Phase between MUA signals is shown for 0.03 and 10 Hz for all pairs of shanks/tetrodes. Unlike phase distribution between population rate and individual neurons, the distribution of phases between MUA signals is unimodal in both infraslow and high frequencies (cf. Figure 4c-d).

**Figure S6.**
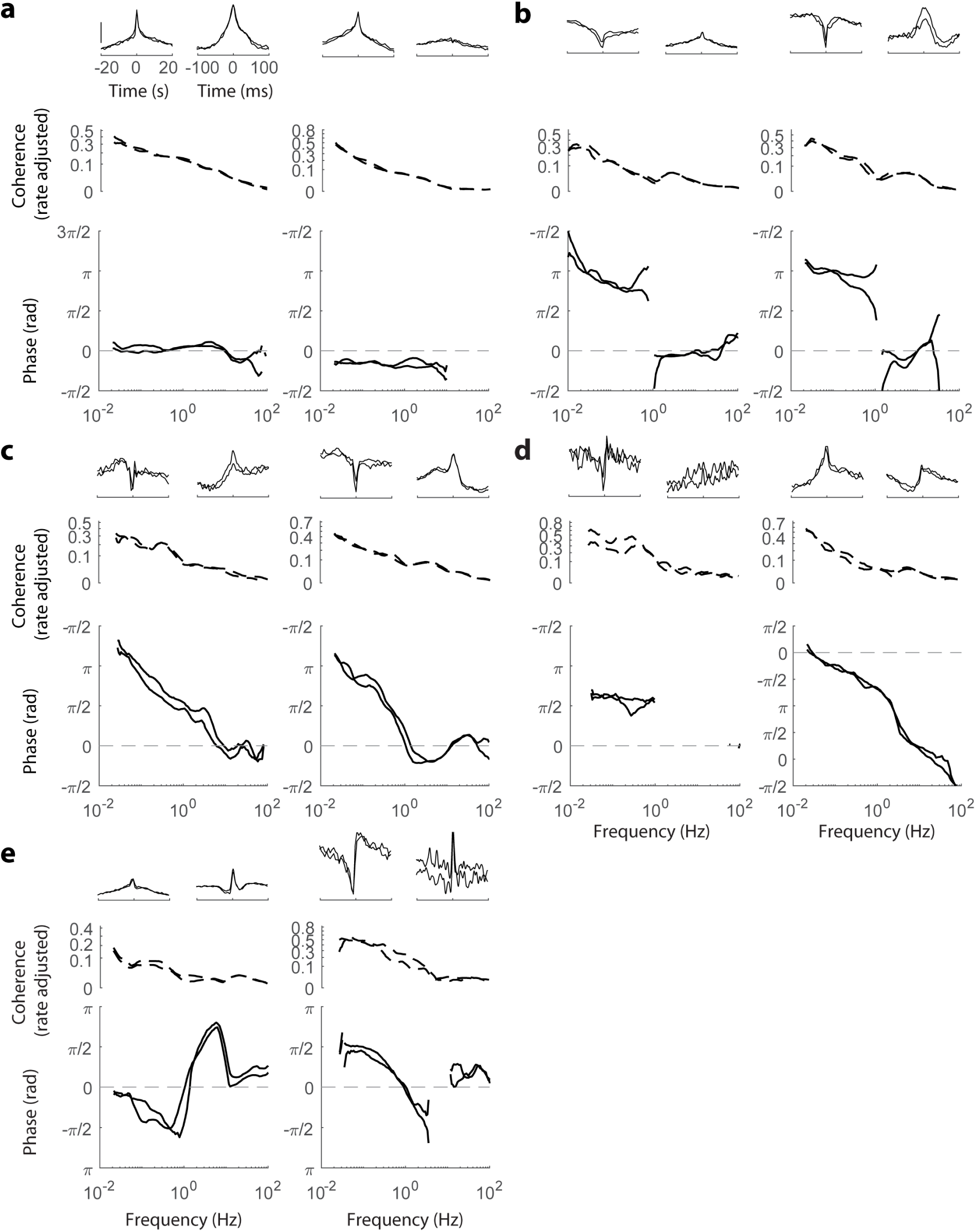
Population coupling phase spectrum. For each example neuron in Figure 5, time domain correlation between the neuron and population rate on fast and slow timescale (top), and its coherence (middle) and phase (bottom) with respect to population rate were evaluated in each half of the recording separately. The values in the two halves closely overlap.

**Figure S7.**
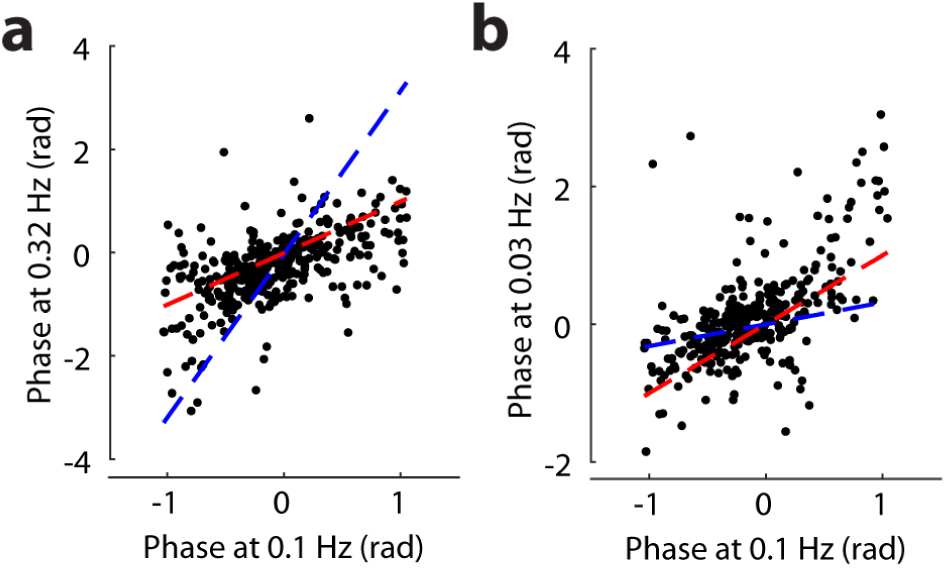
Comparison of linear phase and constant phase models. **(a)** Phase at 0.1 Hz vs phase at 0.32 Hz. Red dashed line indicates identity, corresponding to constant phase model, with R^2^ = 0.23. Linear phase model is shown by blue dashed line, with R^2^ < 0. **(b)** Phase at 0.1 Hz vs phase at 0.03Hz. The dashed lines show the two models as in a. For constant phase model R^2^ = 0.28, for linear phase model R^2^ = 0.17. In a, b the comparison was limited to neurons whose phase at 0.1 Hz was sufficiently close to 0 (specifically, within 1 rad; using other intervals produced similar results).

**Figure S8.**
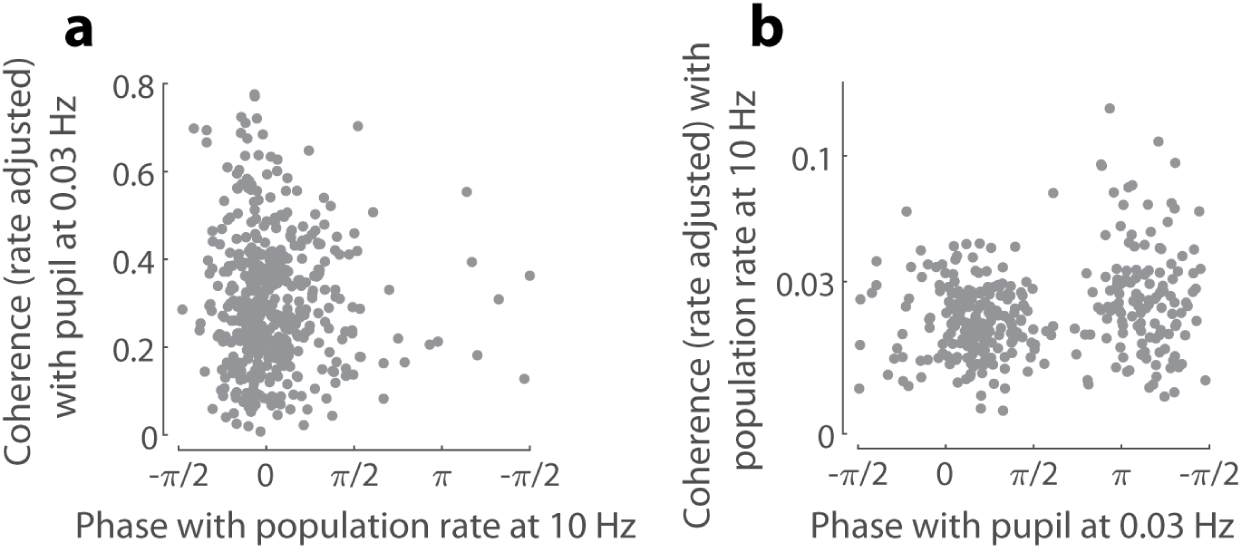
Infraslow pupil coupling vs fast population rate coupling of individual neurons. **(a)** No significant relationship between fast timescale phase and coherence with pupil (P = 0.77). **(b)** A significant correlation between phase with pupil at 0.03 Hz and coherence with population rate at 10 Hz (P < 0.001), where neurons anti-correlated with the pupil are more coherent with population rate on fast timescales. This might be due to sub-classes of cortical neurons differing in their fast and slow timescale population coupling properties.

**Figure S9.**
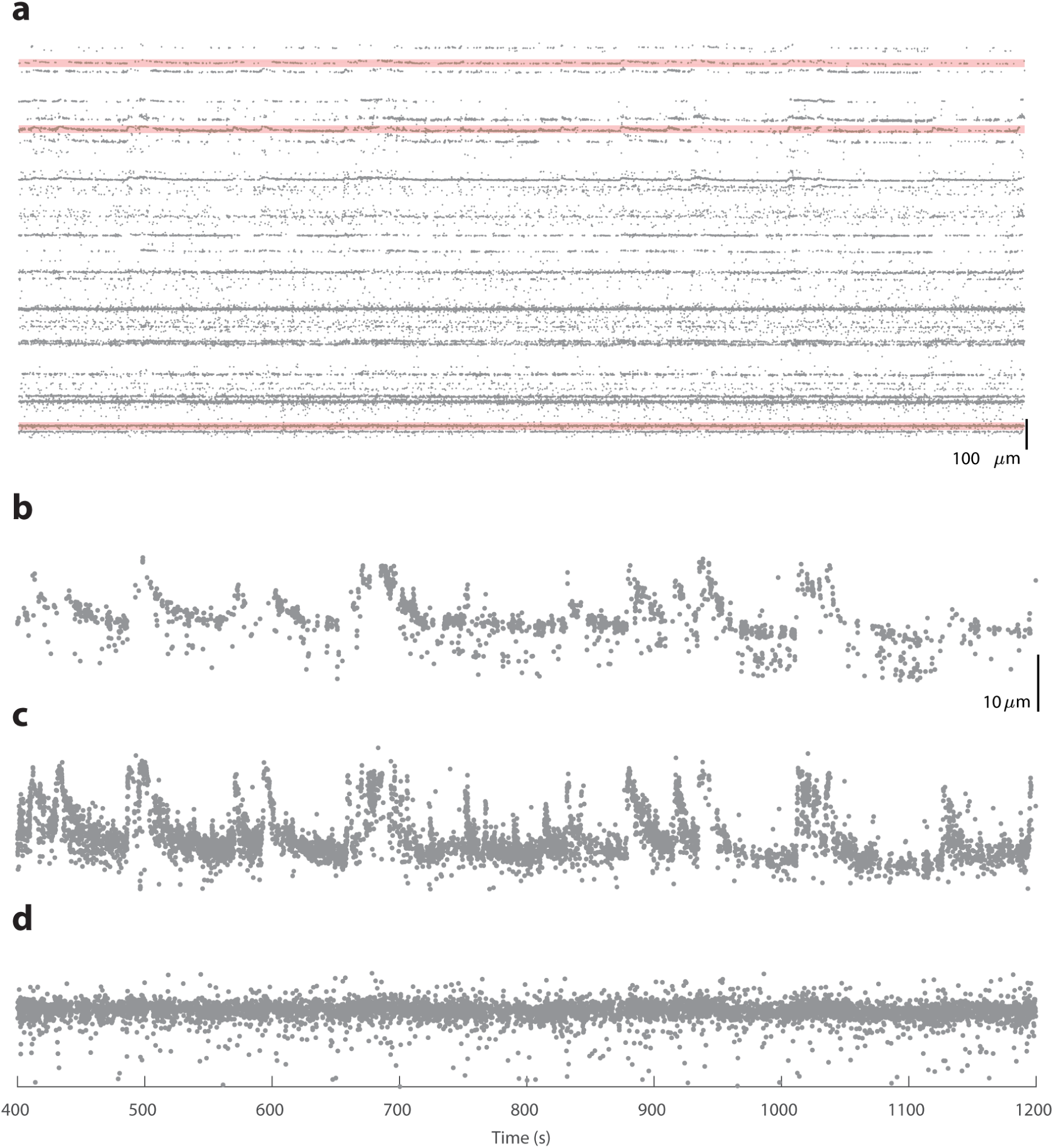
Drift detection in Neuropixels probe recordings. **(a)** Vertical position of each high-amplitude spike detected over 800 s in the top 1.4mm of a Neurpixels probe (the figure looks similar for the rest of the recording, which was omitted) in an example recording, **(b-d)** Three locations on the probe (highlighted in **a),** shown with a higher spatial resolution. In **b, c** drifts of 10-15 μm are clearly visible. The drift events are tens of seconds in duration and occur simultaneously at both locations. Drifts are not present at the third location shown in **d,** ∼1 mm further down the probe.

